# Zinc supplement augments the suppressive effects of repurposed drugs of NF-kappa B inhibitor on ACE2 expression in human lung cell lines in vitro

**DOI:** 10.1101/2021.01.27.428372

**Authors:** Ming-Cheng Lee, Yin-Kai Chen, Yih-Jen Hsu, Bor-Ru Lin

**Affiliations:** Department of Internal Medicine, National Taiwan University Hospital and National Taiwan University College of Medicine, Taipei 10051, Taiwan, R.O.C.; Graduate Institute of Clinical Medicine, College of Medicine, National Taiwan University, Taipei 10002, Taiwan, R.O.C.; Department of Hematology, National Taiwan University Cancer Center, Taipei 10672, Taiwan, R.O.C.; Department of Integrated Diagnostics and Therapeutics, National Taiwan University Hospital, Taipei 10051, Taiwan, R.O.C.

**Keywords:** SARS-CoV-2, COVID-19, ACE2, NF-kappa B, ROS, triclabendazole, emetine, zinc

## Abstract

Severe acute respiratory syndrome coronavirus 2 (SARS-CoV-2) causes a vast number of infections and fatalities worldwide. As the development and safety validation of effective vaccines are ongoing, drug repurposing is most efficient approach to search FDA approved agents against coronavirus disease 2019 (COVID-19). In the present study, we found that endogenous ACE2 expressions could be detected in H322M and Calu-3 cell lines, as well as their ACE2 mRNA and protein expressions were suppressed by pyrrolidine dithiocarbamate (PDTC), a NF-kappa B inhibitor, in dose- and time-dependent manners. Moreover, N-acetyl-cysteine (NAC) pretreatment reversed PDTC-induced ACE2 suppression, as well as the combined treatment of hydrogen peroxide and knockdown of p50 subunit of NF-kappa B by siRNA reduced ACE2 expression in H322M cells. In addition, anthelmintic drug triclabendazole and antiprotozoal drug emetine, repurposed drugs of NF-kappa B inhibitor, also inhibited ACE2 mRNA and protein expressions in H322M cells. Moreover, zinc supplement augmented the suppressive effects of triclabendazole and emetine on ACE2 suppression in H322M and Calu-3 cells. Taken together, these results indicate that ACE2 expression is modulated by reactive oxygen species (ROS) and NF-kappa B signal in human lung cell lines, and zinc combination with triclabendazole or emetine has the clinical potential for the prevention and treatment of COVID-19.

## Introduction

Severe acute respiratory syndrome coronavirus 2 (SARS-CoV-2) caused the coronavirus disease 2019 (COVID-19) pandemic, and more than 95 million confirmed cases were identified worldwide by the beginning of 2021 (COVID-19 Coronavirus Pandemic Update Worldometer). The clinical spectrum of COVID-19 can be very heterogenous, ranging from asymptomatic forms to severe manifestations, including the acute respiratory distress syndrome (ARDS) (*1*). Severe COVID-19 complications are associated with the cytokine storm syndrome, an uncontrolled over-production of inflammation factors which leads to sustain an aberrant systemic inflammatory response contributing to the high mortality rate of the disease (*2*). SARS-CoV-2 already caused more than 2 million fatalities until now, and since there is no specific treatment prescribed to treat COVID-19, therefore, to redevelop an existing drug for licensed use in COVID-19 therapy seems valuable.

Angiotensin-converting enzyme 2 (ACE2) has a multiplicity of physiological roles in renin-angiotensin system (RAS) and amino acid transport. ACE2 receptors are ubiquitous and widely expressed in the heart, vessels, gut, lung, kidney, testis and brain (*3*). At the initial step of SARS-CoV-2 infection, the viral spike (S) glycoproteins interact first with ACE2 receptors on the target cell surface, and are then proteolytically cleaved by trans membrane protease serine 2 (TMPRSS2) to trigger fusion of the viral envelope with the target cell membrane (*4, 5*). Therefore, targeting the Spike-ACE2 binding event has been logically viewed as a promising strategy to mitigate viral infection and control the infection. To date, decoy ACE2 soluble protein (*6*), ACE2 inhibitor and TMPRSS2 inhibitor (*4*) had been investigated for trapping the virus and blocking the binding of spike protein for reducing the accessibility of the virus to host cells (*7*). Their clinical efficacy and safety for COVID-19 patients are still exploring. Therefore, new therapeutic options against this viral entry are urgently desired.

Viral spike protein or dsRNA of SARS-CoV-2 may activate the NF-κB pathway in non-immune cells such as alveolar epithelial cells and endothelial cells, eventually lead to cytokine storm in severe COVID-19 patients (*8*). Therefore, inhibition of NF-κB pathway has been considered to be a potential therapeutic strategy for alleviating the severe form of COVID-19. In patients hospitalized with Covid-19, the use of dexamethasone, acts as NF-kB inhibitor, resulted in lower 28-day mortality among those who were receiving either invasive mechanical ventilation or oxygen alone at randomization but not among those receiving no respiratory support (*9*). The results suggest that the beneficial role of dexamethasone may be at least partly related to the inhibition of NF-κB activation in the severe COVID-19 patients. Abnormal NF-κB signaling is also linked with certain neoplasia, Miller et al had used NF-κB mediated β-lactamase reporter gene assay to screen approximately 2,800 clinically approved drugs, and identified nineteen drugs that have previously unappreciated effects on NF-κB signaling, which may contribute to anticancer therapeutic effects (*10*). Of these, antiprotozoal drug emetine and anthelmintic drug triclabendazole can inhibit NF-κB signaling via inhibition of IκBα phosphorylation. Since antiprotozoal and anthelmintic drugs has lower cellular toxicity than those of anti-cancer drugs, emetine and triclabendazole were selected to investigate their effect on ACE2 expression in this study.

Zinc has been shown to mediate antiviral effects through improving the mucociliary clearance of virus, strengthening the integrity of the epithelium, decreasing viral replication, preserving antiviral immunity, attenuating the risk of hyper-inflammation, supporting anti-oxidative effects (*11*). Observational studies have reported that lower baseline zinc levels in hospitalized adults were associated with a higher risk of mortality, complications, and longer hospital stay following SARS-CoV-2 infection (*12*), suggesting potential benefits of zinc supplementation in reducing COVID-19 morbidity and mortality. Moreover, the clinical trials on zinc supplementation in combination with other drugs such as hydroxychloroquine or chloroquine have been registered (*13*), and the anticipated results coming soon. Because zinc supplementation is a cost-efficient, globally available and simple to use option with little to no side effects, it has most potential to in combination with other repurposed drugs against COVID-19 treatment.

ACE2 expression in Huh7 (hepatocellular carcinoma cells) is increased by the treatment of AMPK activator, through NAD-dependent deacetylase SIRT1 (silent information regulator T1) dependent pathway. (*14*). However, the detail mechanism of ACE2 expression in lung cells is still remain unclear. In this present study, we found ACE2 expressions in lung cancer cells were suppressed by NF-κB inhibitor, ammonium pyrrolidine dithiocarbamate (PDTC), and clinical drugs, triclabendazole and emetine, which are able to inhibit NF-kB signaling. Moreover, zinc supplement augmented the suppressive effects of triclabendazole and emetine on ACE2 expression in lung cancer cells. These findings anticipate that the combination of repurposed inhibitor drugs of NF-κB and zinc supplementary may have a potential to alleviate and prevent the severe COVID-19 pandemics.

## Results

### NF-κB inhibitor, PDTC, decreases endogenous ACE2 expressions in H322M and Calu-3 lung cancer cells

SARS-CoV-2 spike protein binds cell surface ACE2 proteins of target cells to initiate viral entry, agents possessing the interaction blocking could be possible therapeutic strategies for the prevention and treatment of SARS-CoV-2 infection. In order to explore the regulatory mechanism of ACE2 expressions in lung cells, the expression levels of endogenous ACE2 protein were evaluated in 6 lung cancer cell lines by western blot assay. It was found that the highest and middle expression levels of ACE2 were detected in Calu-3 and H322M cells, respectively, compared to the other 4 cell lines (Fig. 1A). Therefore, Calu-3 and H322M cells were selected for the following experiments. Inhibition of NF-κB signaling has a potential therapeutic role in alleviating the severe form of COVID-19 through reduction in the cytokine storm (*15*). To further explore whether NF-κB pathway also play a role in ACE2 regulation, PDTC, an inhibitor of NF-κB activation, was used to treat cells and the cytotoxic effect of PDTC was investigated by MTS assay. After PDTC treatment, we found that 200 μM PDTC did not lead to critical cell death (Fig. 1B), therefore. this concentration was used to process for following experiments. To explore the PDTC effect on mRNA expression of *ACE2*, both H322M and Calu-3 cells were treated with a series dose of PDTC and the mRNA expressions were evaluated. The results showed that PDTC treatment decreased the RNA levels of *ACE2* in a dose-dependent manner in these two cell lines (Fig. 1C and 1D). In addition, these two cell lines were also treated PDTC, and the protein expressions of ACE2 were examined at different time points. The results showed that PDTC treatment obviously decreased the ACE2 protein expressions in a time-dependent manner in H322M and Calu-3 cells (Fig. 1E and 1F). These findings indicate that ACE2 expression is regulated by NF-κB signal pathway, and implicate that clinical drugs with the activity of NF-κB inhibition may exert beneficial effects against covid-19 disease.

**Figure 1.**
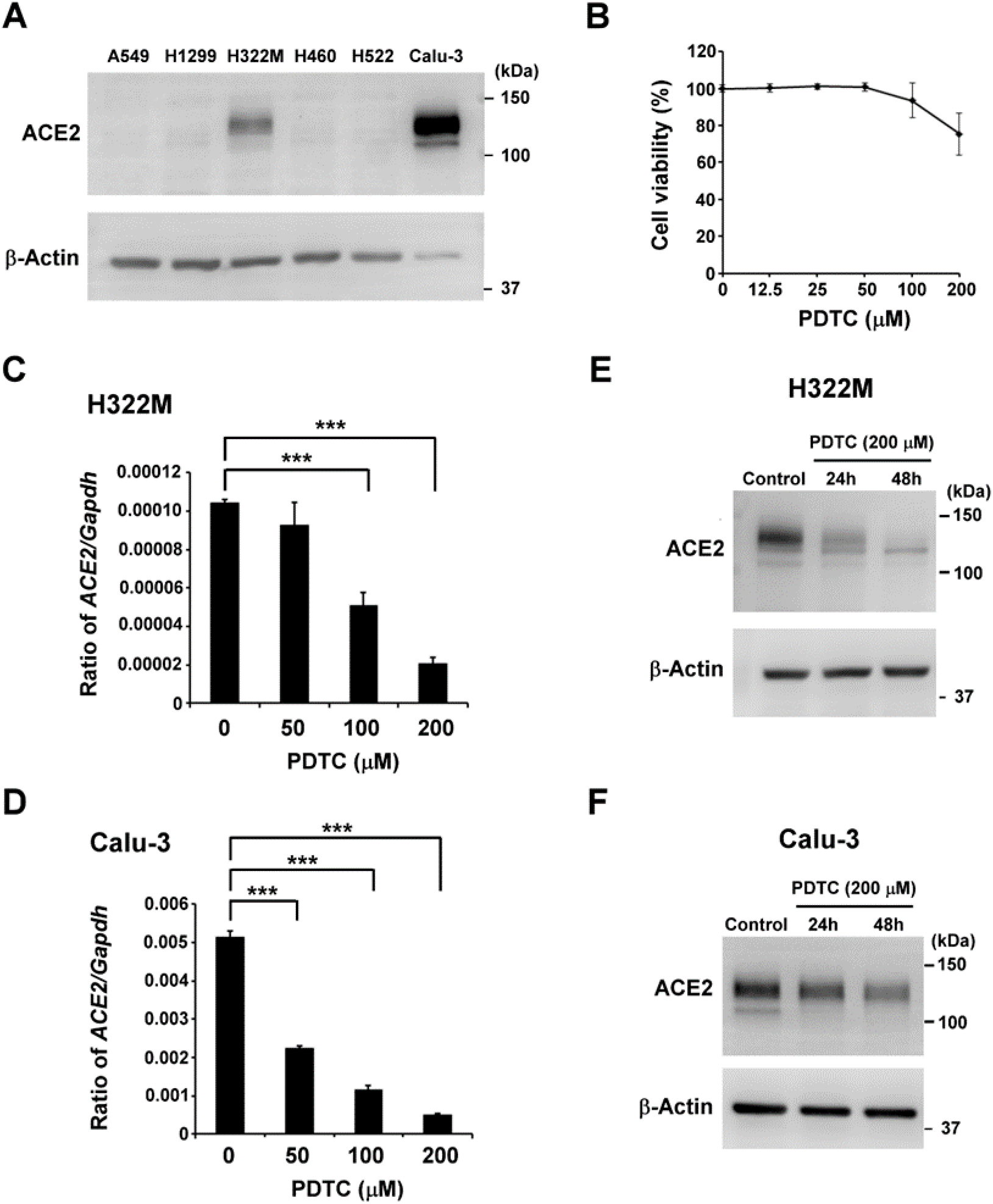
NF-κB inhibitor treatment inhibiteds endogenous ACE2 expressions of H322M and Calu-3 lung cancer cells. (A) Endogenous ACE2 protein expressions of 6 different lung cancer cells were investigated by Western blot assay using β-actin as loading control. (B) H322M cells were seeded in 96-well plates and treated with a concentration series of PDTC (0, 12.5, 25, 50, 100 or 200 μM) for 48 hours, then the relative cell viabilities were determined by MTS assay. Error bars represented the standard deviation. The viability of control cells was designated as 100%, and that of the other groups were expressed as percent of the control. (C and D) H322M and Calu-3 cells were treated with a concentration series of PDTC (0, 50, 100 or 200 μM) for 24 hours and the total RNAs were collected for *ACE2* measurement by real-time qRT-PCR assays. The data were normalized by GAPDH expression and presented as mean ± SEM (n = 3). (***p < 0.0005). (E and F) H322M and Calu-3 cells were treated with 200 μM PDTC, then harvested at different time points after treatment. Treated cells were lysed for Western blot analysis of ACE2, and β-actin was used as loading control.

### N-acetyl-cysteine blocks PDTC-derived ACE2 suppression in H322M cells

To further explore the underlying mechanism of PDTC-derived ACE2 suppression, p50 subunit of NF-κB was inhibited by its specific siRNA and its associated effect on ACE2 protein expression were evaluated. It was found that p50 proteins were obviously inhibited by the specific siRNA, however, ACE2 protein expression was not obviously affected in H322M cells (Fig. 2A). Previous studies had indicated that PDTC is a metal-chelating compound that exerts both pro-oxidant and antioxidant effects (*16*). To explore whether ROS involves in PDTC-derived ACE2 suppression, H322m cells were pretreated with N-acetyl-cysteine (NAC) to inhibit ROS production during PDTC treatment. We noted that NAC treatment alone enhanced ACE2 mRNA and protein expressions, whereas NAC pretreatment blocked PDTC-induced ACE2 suppression in both of mRNA and protein levels (Fig. 2 B and 2C), suggesting PDTC acts as a pro-oxidant in H322M cells. To mimic the condition caused by PDTC in H322M cells, H322M cells with p50 knockdown were treated with H2O2 and ACE2 expressions were evaluated. Immunoblotting of PARP was conducted to monitor cell death level. It was found that H2O2 treatment suppressed ACE2 expressions in p50-silenced H322M cells, but not in control cells (Fig. 2D). These results together suggest that PDTC suppresses ACE2 expression through ROS induction and NF-κB inhibition in H322M cells.

**Figure 2.**
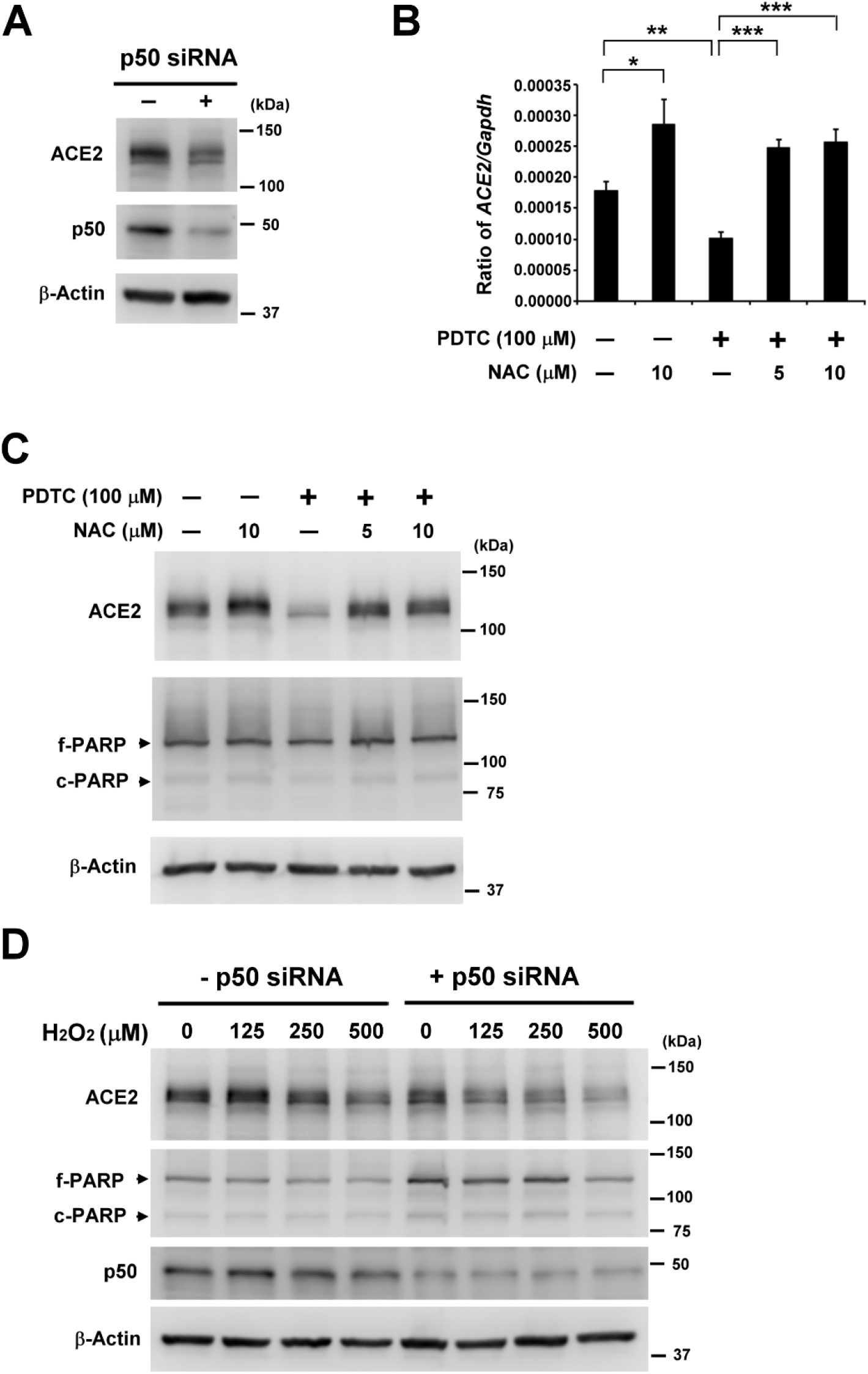
Thiol antioxidant NAC blocks PDTC-derived ACE2 suppression in H322M cells. (A) H322M cells were electroporated with control or p50 siRNA for 48 hours, then cells were lysed for Western blot analysis of ACE2 and p50, meanwhile β-actin was used as loading control. (B) PDTC (100 μM) was used to treated H322M cells with 30 minutes pretreatment of NAC or not. And the total RNAs were collected for *ACE2* measurement by real-time qRT-PCR assays. The data were normalized by GAPDH expression and presented as mean ± SEM (n = 3). (*p < 0.05, ** p < 0.005, and ***p < 0.0005). (C) As 2B conditions, the treated cells were lysed for Western blot analysis of ACE2 and PARP, meanwhile β-actin was used as loading control. The position of the native PARP (f-PARP) and the cleaved fragment (c-PARP) is indicated. (D) H322M cells were electroporated with control or p50 siRNA for 24 hours, then cells were treated with H2O2 for an additional 24 hours. Treated cells were lysed for Western blot analysis of ACE2, p50 and PARP, meanwhile β-actin was used as loading control.

### Triclabendazole or emetine administration suppresses endogenous ACE2 expression of H322M cells

Drug repurposing is a fast approach to using clinical approved drugs for different disease treatments bypassing the long streamline of the drug discovery process. Base on the concept, NF-κB inhibitor, triclabendazole and emetine (*10*), were selected to investigate their effects on ACE2 expression of lung cancer cells. First, the cytotoxic effects of triclabendazole and emetine against H322M cells were investigated to identify the suitable dose range of triclabendazole and emetine for experiments. It was found that 100 μM triclabendazole and 125 nM emetine treatment did not caused apparently inhibition of cell growth (Fig. 3A). To further explore the effects of triclabendazole or emetine on mRNA expression of *ACE2*, H322M cells were treated with different concentrations of triclabendazole or emetine, and we fund that these two drugs suppressed *ACE2* expressions in a dose-dependent manner (Fig. 3B). The treatment of 100 μM triclabendazole or 100 nM emetine individually inhibited a half of *ACE2* mRNA levels of H322M cells. As well as the treatment of triclabendazole or emetine also decreased the ACE2 protein expressions in a time-dependent manner (Fig. 3C). Immunoblotting of PARP was conducted to monitor cell death level, and there was no apparently cleaved PARP, a marker of apoptosis, was detected, except of the cells with triclabendazole treatment for 48 hours. These findings suggest that triclabendazole and emetine, FDA approved clinical drugs, exhibit anti-SARS-CoV-2 activity via ACE2 suppression to reduce viral infection.

**Figure 3.**
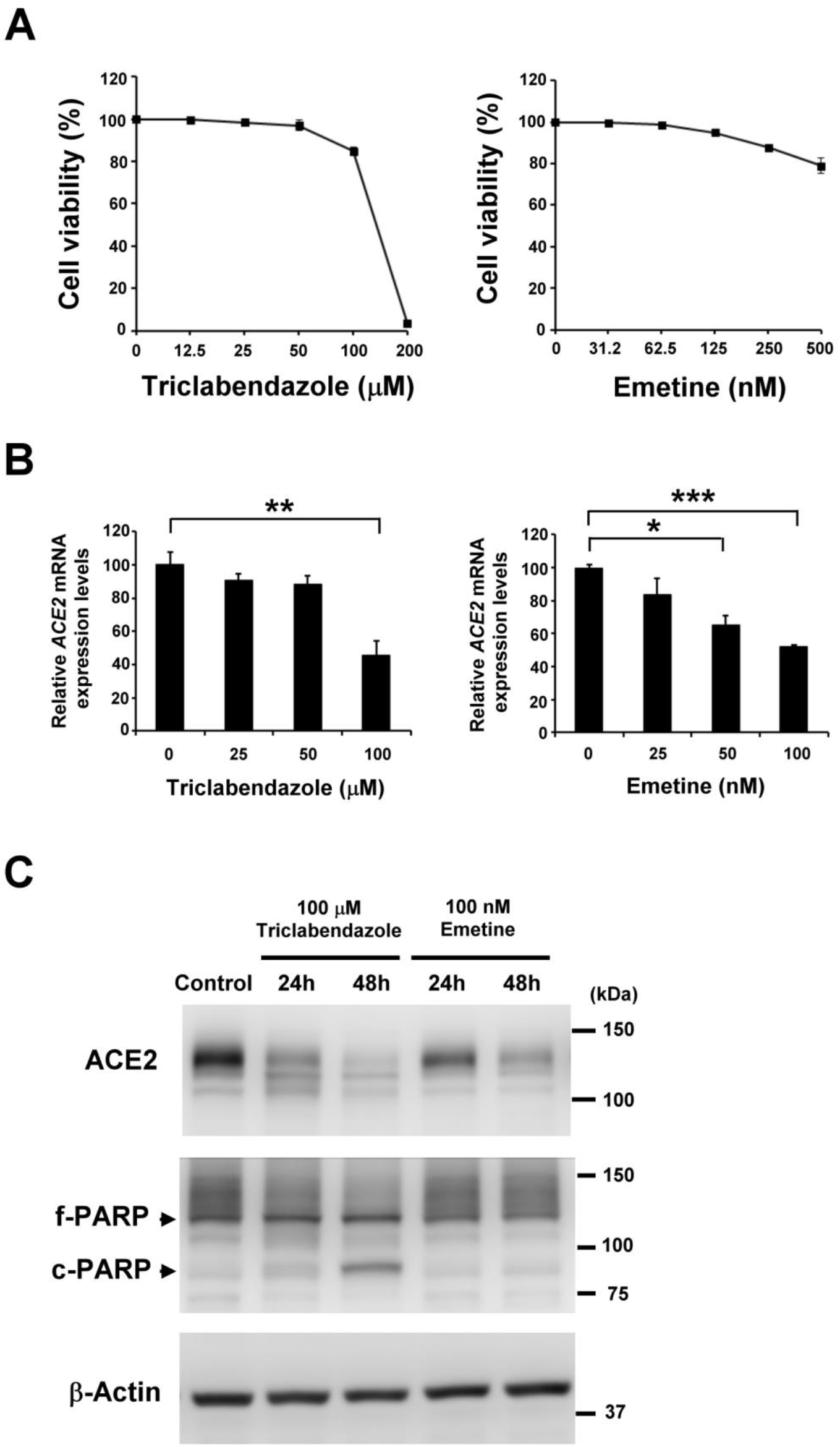
Endogenous ACE2 expressions of H322M cells are suppressed by clinical drugs, triclabendazole and emetine. (A) H322M cells were seeded in 96-well plates and treated with a concentration series of triclabendazole (0, 12.5, 25, 50, 100 or 200 μM) or emetine (0, 31.2, 62.5, 125, 250, or 500 nM) for 48 hours, then the relative cell viabilities were determined by MTS assay. Error bars represented the standard deviation. The viability of control cells was designated as 100%, and that of the other groups were expressed as percent of the control. (B) H322M cells were treated with a concentration series of triclabendazole (0, 50, or 100 μM) or emetine (0, 50, or 100 nM) for 24 hours and the total RNAs were collected for *ACE2* measurement by real-time qRT-PCR assays. The data were normalized by GAPDH expression and presented as mean ± SEM (n = 3). (*p < 0.05, ** p < 0.005, and ***p < 0.0005). (C) H322M cells were treated with 100 μM triclabendazole or 100 nM emetine, then harvested at different time points after treatment. Treated cells were lysed for Western blot analysis of ACE2 and PARP, meanwhile β-actin was used as loading control. The position of the native PARP (f-PARP) and the cleaved fragment (c-PARP) is indicated.

### Zinc supplement augmentes the suppressive effects of triclabendazole and emetine on ACE2 expressions in H322M and Calu-3 cells

Combination therapy is a valuable approach to use different drugs blocking similar pathways to increase treatment efficacy and reduce the dose and side effects of toxic drugs. Zinc is a trace element with potent immunoregulatory and antiviral properties, and is utilizing in the treatment of COVID-19, even its clinical outcome is still unclear (*17*). Here, we investigated the effects of triclabendazole or emetine alone, or its combination with zinc sulfate on ACE2 expression of the lung cell lines. First, it was found that *ACE2* mRNA and protein was slightly repressed by the treatment of 300 μM zinc sulfate (Fig. 4A and 4B). To investigate whether zinc treatment combined with triclabendazole or emetine has the ability to reduce the drug dose for ACE2 suppression, H322M and Calu-3 cells were treated with triclabendazole, emetine, zinc sulfate alone, or in combination with zinc sulfate and each of these two clinical drugs for 24 hours. We proved that ACE2 protein expressions did not be dramatically repressed in H322M and Calu-3 cells treated with triclabendazole, emetine or zinc sulfate alone, whereas zinc treatment combined with triclabendazole or emetine apparently suppressed the ACE2 protein expressions (Fig. 4C and 4D). Meanwhile, there was no apparently cleaved PARP, a marker of apoptosis, was detected in these two cell lines. These findings anticipate that triclabendazole or emetine plus zinc supplementation may be more effective in reducing virus amplification through decreasing ACE2 protein expression than triclabendazole or emetine monotherapy.

**Figure 4.**
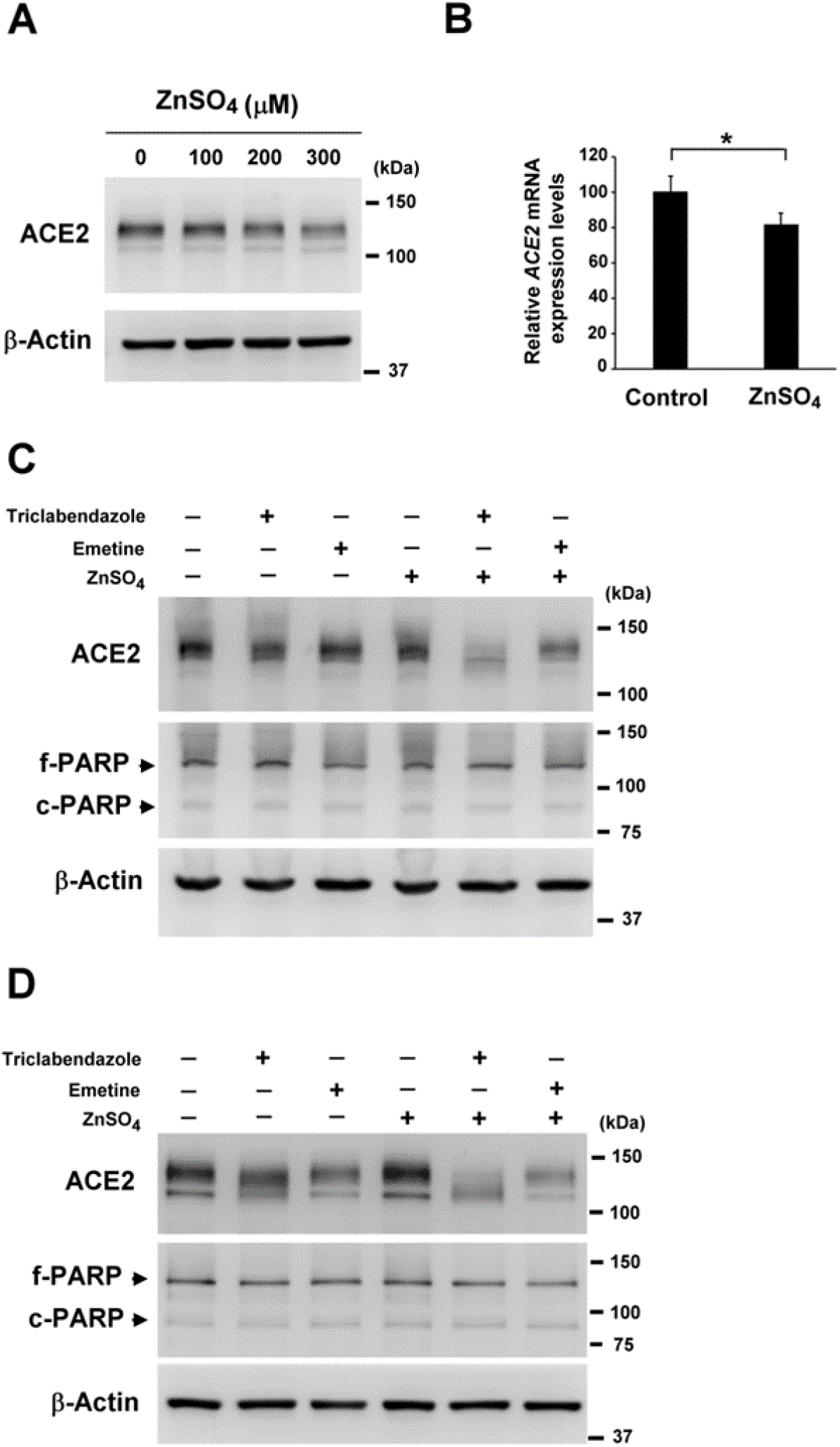
Combinational treatment of zinc sulfate and clinical NF-kB inhibitors synergistically suppresses ACE2 expressions in H322M cells. (A) H322M cells were treated with a concentration series of zinc sulfate (0, 100, 200 or 300 μM) for 24 hours, then the treated cells were lysed for Western blot analysis of ACE2 and PARP, meanwhile β-actin was used as loading control. (B) H322M cells were treated with ZnSO4 (300 μM) for 24 hours and the total RNAs were collected for *ACE2* measurement by real-time qRT-PCR assays. The data were normalized by GAPDH expression and presented as mean ± SEM (n = 3). (*p < 0.05). (C) H322M cells and (D) Calu-3 cells were treated with triclabendazole (50 μM), emetine (100 nM), zinc sulfate (150 μM) alone, or in combination with zinc sulfate (150 μM) and each of these clinical drugs for 24 hours. Treated cells were lysed for Western blot analysis of ACE2 and PARP, meanwhile β-actin was used as loading control. The position of the native PARP (f-PARP) and the cleaved fragment (c-PARP) is indicated.

## Discussion

Globally, a number of repurposing drugs, known to act on interruption of virus life cycle and modulation of host immune response, are now being investigated in clinical trials for the patients with moderate-to-severe COVID-19. For example, remdesivir (an RNA polymerase inhibitor) was able to reduce the time to recovery in adults who were hospitalized with COVID-19 and had evidence of lower respiratory tract infection (*18*). Immunosuppressive drugs, such as tocilizumab (*19*), hydrocortisone (*20*), baricitinib (*21*) and dexamethasone (*22*) have been evaluating in clinical trials, however, until now there is still lack of the overwhelming evidence supporting their beneficial effects on COVID-19 therapy. Therefore, searching for FDA approved drugs that can effectively control SARS-CoV-2 infection are urgently needed. In this present study, we demonstrated that endogenous ACE2 expression could be repressed by NF-κB inhibitor, as well as triclabendazole and emetine, act as immunosuppressive drugs through NF-κB inhibition, also exhibit the ability to suppress ACE2 expression of human lung cell lines, implicating their potential to mitigate the virus’s accessibility.

As the respiratory droplets carrying infectious SARS-Cov-2 virus are inhaled into the upper air way, viral spike (S) protein will preferentially bind to the membrane protein ACE2 on epithelial cells for virus entry. Moreover, previous study in patients with COVID-19 indicated that airway epithelial cells showed an average three-fold increase in expression of the SARS-CoV-2 entry receptor ACE2 (*23*), suggesting that ACE2 increase induced by virus infection may facilitate viral amplification. Therefore, to understand the factors affecting ACE2 expression may help us to prevent ACE2 induction during SARS-CoV-2 virus infection. We had explored the endogenous ACE2 expressions in six different lung cell lines and found that the highest and middle levels of ACE2 expressions were detected in Calu-3 and H322M cells, respectively (Fig. 1A). Calu-3 has the highest expression of ACE2 may be due to Calu-3 is a mucus-producing submucosal gland carcinoma cell line derived from human bronchial epithelium (*24*), which is consistent with that the highest expression of *ACE2* was found in nasal secretory cells (*25*).

COVID-19 patients with severe COVID-19 displayed the uncontrolled NF-κB-related cytokine release syndrome (CRS), which is responsible for the poor prognosis and multiorgan defects (*8*). Thus, the pharmacological blockade of the NF-κB signaling pathway can strongly represent a potential therapeutic consideration to attenuate COVID-19. For example, hydrocortisone and dexamethasone were proved to exhibit the activity of NF-κB inhibition, meanwhile their clinical trials for Covid-19 therapy are ongoing (*26, 27*). Here, we proved that administration of pyrrolidine dithiocarbamate (PDTC) inhibited the endogenous *ACE2* mRNA and protein expressions in Calu-3 and H322M cells in a dose- and time-dependent manner (Fig. 1C-1F), suggesting that activation of NF-κB signaling is required for ACE2 expression in lung epithelial cells. While blocking NF-κB signaling via p50 siRNA, however, ACE2 expression was not dramatically repressed in H322M cells (Fig. 2A), indicating that NF-κB inhibition and a PDTC-derived another event cooperate in PDTC-induced ACE2 suppression in H322M cells. Previous study had indicated that PDTC is a metal-chelating compound that exerts both pro-oxidant and antioxidant effects (*16*). In addition, thiol antioxidants reversed PDTC actions on blocking DNA binding activity of NF-κB (*28*). Here, we found that the pretreatment of thiol oxidant, N-acetyl-cysteine (NAC), abolished PDTC-induced ACE2 suppression in both of mRNA and protein levels (Fig. 2B and 2C). Moreover, the administration of hydrogen peroxide augmented the inhibition of ACE2 expression in H322M cells with p50 silenced (Figure 2D), implicating the pro-oxidant role of PDTC involves in ACE2 suppression. More studies need to be done for dissecting of detailed mechanism of PDTC-induced ACE2 suppression in the future.

To explore the potential clinical application of our findings, we sought FDA-approved drugs with the property of NF-κB inhibition to explore their effects on ACE2 expression in H322M cells. In the present study, antiprotozoal drug emetine and anthelmintic drug triclabendazole were selected to investigate their effects on ACE2 expression. The IC50 values of inhibiting IκBα phosphorylation for emetine and triclabendazole are 0.31 μM and 25.1 μM, respectively (*10*). Administration of 100 nM emetine or 100 μM triclabendazole significantly repressed ACE2 expressions in both of mRNA and protein levels in H322M cells (Fig. 3B and 3C). It is noteworthy that the serum levels of inflammatory cytokines were significantly reduced by emetine treatment in the pulmonary hypertension rat model (*29*). Moreover, triclabendazole treatment obviously reduced inflammatory cytokine levels, indicating the improvement of the immunological responses of the patients with acute or chronic stage of fasciolosis (*30*). The results in the present and earlier studies suggest that emetine and triclabendazole may exhibit therapeutic potential in COVID-19 patients through breaking of virus life cycle and modulation of host immune response.

Combination therapy has proven successful in a variety of medical areas, such as infectious and metabolic diseases. The advantages of combination therapy include to increase treatment efficacy, to reduce the dose and side effects of toxic drugs, prevent the development of drug resistance, and to reduce the duration of treatment. Previous studies had implicated that zinc may possess antiviral activity and anti-inflammatory activity through inhibition of SARS-CoV RNA polymerase and disruption of NF-κB signaling, respectively (*17, 31*). In addition, it was noted that 57.4% of the COVID-19 patients were zinc deficient, as well as these patients had higher rates of complications, acute respiratory distress syndrome, prolonged hospital stay, and increased mortality (*12*). Despite the lack of clinical data, the previous findings suggest that modulation of zinc status may be beneficial in COVID-19. In this study, we found that zinc administration alone did not apparently affect *ACE2* mRNA and protein levels in H322M cells, exception of high dose treatment (Fig. 4A and 4B). By contrast, the apparent ACE2 suppressions were observed 24 hours only after combined treatment of 150 μM zinc sulfate with 50 μM triclabendazole or 100 nM emetine in H322M and Calu-3 cells (Fig. 4C and 4D), compared with the monotreatment of triclabendazole or emetine. Taken together, these findings indicate that zinc supplement exhibits the ability to augment the inhibitive effect of triclabendazole and emetine on ACE2 suppression, and implicate that zine supplement may increase the potential clinical application of triclabendazole or emetine for COVID -19 prevention and treatment.

Besides immunosuppressive functions of zinc, triclabendazole and emetine, we demonstrated that zinc administration also enhanced the suppressive effects of triclabendazole and emetine on endogenous ACE2 expressions of H322M and Calu-3 cells, hypothesizing that triclabendazole or emetine plus zinc supplementation may be more effective in reducing COVID-19 morbidity and mortality than triclabendazole or emetine monotherapy. Therefore, triclabendazole or emetine in combination with zinc should be considered as additional study arm for COVID-19 clinical trials. Given ACE2 plays many important biological roles in regulating cardiovascular, renal and innate immune systems, therefore, leveraging ACE2 as a therapeutic target for COVID-19 requires collective considerations to balance the pros and cons. In order to maximize therapeutic effectiveness while minimizing collateral adverse effects, inhalable delivery would seem to be a logical option to selectively and locally deposit triclabendazole or emetine combined with zinc to lung epithelial cells in the airway and airspace compartments. To prove the hypothesis, of cause, more works have to be done.

## Materials and Methods

### Reagents

PDTC (1-Pyrrolidinecarbodithioic Acid, Ammonium Salt, 548000; merckmillipore), triclabendazole (HY-B0621; MCE) and emetine dihydrochloride (HY-B1479A, MCE) were dissolved in dimethyl sulfoxide (DMSO), whereas, ZnSO4 (Zinc sulfate heptahydrate, Z0501; Sigma-Aldrich) and NAC (N-Acetyl-L-cysteine, A9165; Sigma-Aldrich) were dissolved in ddH2O. The solvent was routinely used in the control group of experiment.

### Cell culture

Lung cancer cells A549, H322M, H460 and H522 were cultured in Roswell Park Memorial Institute (RPMI) 1640, and H1299 cells were cultured in Dulbecco’s modified Eagle’s medium (DMEM). Calu-3 cells were courtesy of Dr. Tsung-Tao Huang (National Applied Research Laboratories, Hsinchu, Taiwan) and were cultured in Minimum Essential Medium (α-MEM) (Gibco BRL, Grand Island, NY, USA). All culture media were supplemented with 10 % heat-inactivated fetal bovine serum, 1 % penicillin/streptomycin solution (Gibco-BRL). The cells were grown in a humidified incubator containing 5% CO2 at 37 °C.

### MTS assay

To determine the cytotoxicity of PDTC, triclabendazole and emetine for H322M, cells were seeded in 96-well plates overnight and treated for 48 hours with different concentrations of PDTC (12.5-200 μM) or triclabendazole (12.5-200 μM), or emetine (31.2-500 nM) or DMSO as a vehicle control. The cell viability was then evaluated by MTS assay. Twenty microliters of the MTS CellTiter 96 Cell Proliferation Assay reagent (Promega Corporation, Madison, WI, USA) were added to each well. After 1 hours of incubation at 37°C, the 490 nm absorbance of colored product was measured in a EPOCH2 microplate reader (BioTek Instruments, Inc., USA).

### Real-time quantitative RT-PCR (qRT-PCR) analysis

To quantify the endogenous mRNA expression levels, real-time quantitative RT-PCR (qRT-PCR) was performed. Total RNAs of cells were isolated using PureLink® RNA Mini Kit (ambion by Life technologies, Carlsbad, CA, USA) according to the manufacturer’s instructions. One microgram of total RNA was used to synthesize the cDNA using IQ2 MMLV RT-Script (Bio-Genesis) with Oligo-dT primer (Gene Teks). Quantitative PCR was performed using Kapa SYBR fast qPCR master mix (Kapa Biosystems, Woburn, MA, USA). The primer sequences for qPCR are GAPDH-fw: 5’-GAAGGTGAAGGTCGGAGT-3’ and GAPDH-rev: 5’-GAAGATGGTGATGGGATTTC-3’; ACE2-fw: 5’-GGACCCAGGAAATGTTCAGA-3’ and ACE2-rev: 5’-GGCTGCAGAAAGTGACATGA-3’. The qPCR reaction was run in a final volume of 20 μL containing 1 μL of reverse transcriptase product, 10 μL of 2X Kapa SYBR fast qPCR master mix, and 0.6 μL of each primer (10 μM). The qPCR mixtures were pre-incubated at 95°C for 3 min, followed by 40 cycles of 95°C for 3 s and 60°C for 30 s. All qPCR was performed in triplicates by using CFX Connect^™^Real-Time System (Bio-Rad Laboratories, Inc.).

### Western blot analysis

To evaluate the protein expression levels after drugs treatment, western blot analysis was performed. The cells were dissolved in a lysis buffer containing 50 mM Tris-HCl, pH 7.4, 150 mM NaCl, 0.25% deoxycholic acid, 1% NP-40, 1 mM EDTA, and protease inhibitor Cocktail Set III (Calbiochem). Fifty micrograms of protein was fractionated via 10% SDS-PAGE and transferred to Immobilon Transfer Membranes (Millipore Corporation, Billerica, MA, USA). Primary antibodies used in this study included those against β-actin (GTX109639; dilution 1:10,000; GeneTex, Inc.), poly (ADP-ribose) polymerase (PARP) (9542S; dilution 1:1,000; Cell Signaling Technology, Inc.), ACE2 (bs-1004R; dilution 1:1,000; Bioss, Inc.), and p50 (Santa Cruz Biotechnology, Inc., USA; dilution 1:1000). The HRP-conjugated secondary antibodies recognizing mouse-IgG (7076) or rabbit-IgG (7074) were purchased from Cell Signaling Technology, Inc. (dilution 1:5000∼10000). The chemiluminescence signal was recorded using the BioSpectrum UVP810 Imaging System with VisionWorks software (GE Healthcare Life Sciences).

### Knockdown of p50 by electroporation

To knock down p50 RNA expression, scrambled control and SMARTpool siRNA targeting p50 (Dharmacon, Lafayette, CO, USA) were delivered into H322M cells using Neon transfection system (Invitrogen) according to the manufacturer’s instructions. Forty-eight hours after incubation in cell culture, the knockdown efficiencies were determined by protein levels of the targeted genes using Western blot assay.

### Statistical analysis

All values were expressed as mean ± standard error. The Student’s t-test was used to analyze the results, and a P value of less than 0.05 was considered statistically significant.

## Acknowledgments

We thank the staff of the Second Core Lab, Department of Medical Research, National Taiwan University Hospital for technical support during the study.

## Authors Contributions

Conceptualization, M.C.L. and B.R.L.; Methodology, M.C.L. and Y.J.H.; Validation, M.C.L., Y.K.C. and B.R.L.; Writing – Original Draft Preparation, M.C.L.; Writing – Review & Editing, Y.K.C. and B.R.L.

